# A ligand-receptor interactome atlas of the zebrafish

**DOI:** 10.1101/2022.12.15.520415

**Authors:** Milosz Chodkowski, Andrzej Zieleziński, Savani Anbalagan

**Affiliations:** Institute of Molecular Biology and Biotechnology, Faculty of Biology, Adam Mickiewicz University in Poznań, Poznań, Poland

**Keywords:** Secretome, Receptome, Interactome, synapse, PSD, glia, pituicyte, oxytocin, Zebrafish

## Abstract

Studies in zebrafish can unravel the functions of cellular communication and thus identify novel bench-to-bedside drugs targeting cellular communication signaling molecules. Due to the incomplete annotation of zebrafish proteome, the knowledge of zebrafish receptome, secretome and tools to explore their interactome is limited. To address this gap, we *de novo* predicted the cellular localization of zebrafish reference proteome using deep learning algorithm. We combined the predicted and existing annotations on cellular localization of zebrafish proteins, and created repositories of zebrafish secretome, receptome, and interactome as well as associated diseases and targeting drugs. Unlike other tools, our interactome atlas is primarily based on physical interaction data of zebrafish proteome. The resources are available as R and Python scripts (*https://github.com/DanioTalk*). *DanioTalk* provides a novel resource for researchers interested in targeting cellular communication in zebrafish, as we demonstrate in applications studying synapse and axo-glial interactome.

## Introduction

Ligand-receptor-mediated cellular communication is essential for diverse processes such as differentiation, homeostasis, tissue morphogenesis, animal development, and behaviour^1–4^. Dysregulated cellular communication can lead to a plethora of phenotypes ranging from uncontrolled cellular proliferation, developmental defects to disease-states and drug resistance^5–7^. Hence, receptors and ligands are among the leading drug targets for treating human disease conditions^8,9^.

With advances in cryo-electron microscopy and machine learning approaches that enable protein structure prediction, the number of drug targets is expected to increase^10–12^. These developments can aid drug screenings directly using animal model organisms. Due to challenges in using mammalian models in drug screening and limitations of *in vitro* drug screens, larval zebrafish are being increasingly used in high throughput drug screening^13–16^. However, despite the importance of zebrafish in drug discovery, there are no actively maintained datasets of zebrafish secreted factors, receptors and their interactions. Few existing predictions of secretome and membrane proteome are based on incomplete annotation data^17–20^.

With the extensive single-cell RNA-seq datasets of zebrafish, cellular communication in zebrafish can be explored at a holistic level even for rare and transient cells^21,22^. Several tools built on mammalian interactome records have been applied to map the zebrafish ligand-receptor interactions ^23–25^. However, only about half of the zebrafish secretome and receptome data have orthologous interaction records for human genes. Tools that depend on such ortholog records are likely to ignore zebrafish genes lacking mammalian orthologs or genes with incomplete orthologue records. Hence, it is necessary to build a zebrafish secretome and receptome database that also includes information on ortholog-less genes and genes with incomplete annotation data.

Here, we describe *DanioTalk*, a repository of zebrafish secretome, receptome, and interactome and tools to explore them from zebrafish-based omics datasets. To address the limitations of existing tools, *DanioTalk* secretome and receptome datasets are based on deep learning-based subcellular localization predictions allowing the incorporation of orphan zebrafish proteins. *DanioTalk* can identify secretome-receptome pairs primarily based on physical interaction records of zebrafish proteome. *DanioTalk* scripts are available on GitHub and allow exploration of ligand-receptor interactions in any zebrafish-based -omics datasets. Finally, we present the application of *DanioTalk* on mapping the secretome-receptome interactions between synaptosome and postsynaptic density (PSD) and axo-glial interactions between oxytocin neurons and glial pituicytes from RNA-seq datasets. We identified novel ligand-receptor interaction pairs with potential functions in axo-glial interactions. *DanioTalk* provides a novel tool and opportunity for the zebrafish research community to explore ligand-receptor interactions.

## Results

### *De novo* prediction of zebrafish secretome and membrane proteome

A bottleneck in using gene ontology (GO) terms or UniprotKB-based cellular localization records of the zebrafish proteome is the incompleteness of such annotations (**Supplementary Figure S1A**)^26,27^. To address this issue, we used DeepLoc 2.0, a sequence-based deep learning algorithm that can predict cellular localization of proteins with high accuracy ^28^ (**Figure 1A**). We subjected the current zebrafish reference proteome (UP000000437) to DeepLoc 2.0-based prediction ^29^. The majority of proteins were predicted to be cytoplasmic (*n* = 12186; 26%), nuclear (*n* = 7905; 17%) and multilocalizing (*n* = 12211; 26%) (**Figure 1B-C, Supplementary Figure S1B & Supplementary Table 1**). DeepLoc 2.0 predicted that 3699 (8%) and 6481 (14%) of proteins were exclusively extracellular- and cell membrane-localized, respectively (**Figure 1B & Supplementary Table 1**). This corresponds to 2571 (10%) and 4002 (15%) genes that code extracellular and membrane-localized proteins, respectively (**Figure 1B’**). In comparison to previous records, DeepLoc 2.0 identified an additional 1104 and 1953 genes coding for extracellular and membranal proteins, respectively (**Figure 1C & D**) ^18,20,27^.

**Figure 1.**
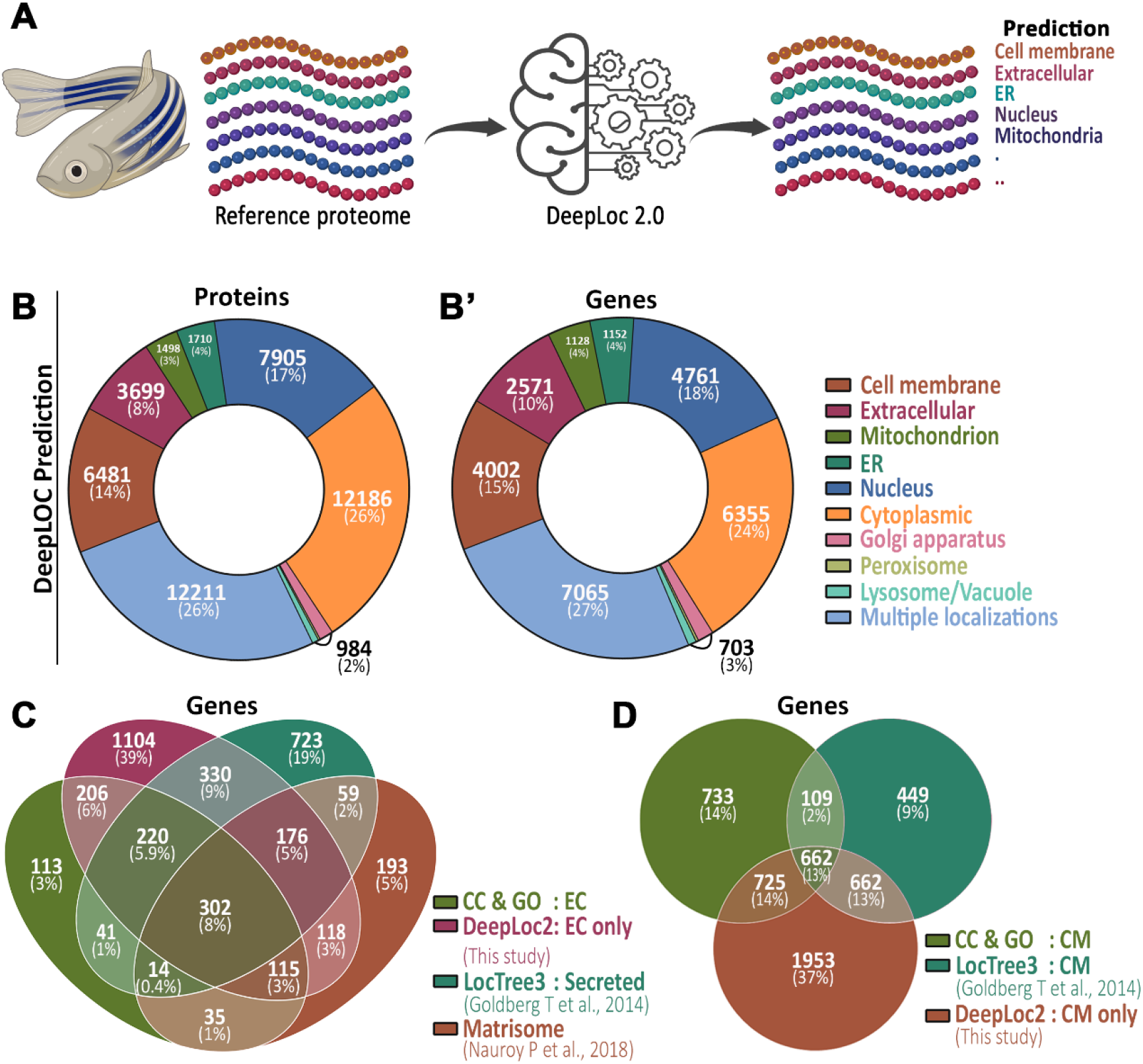
Prediction of cellular localization of zebrafish reference proteome. (A) Graphic describing the DeepLoc 2.0-based prediction of cellular location of the reference zebrafish proteome. (**B**) Zebrafish reference proteome cellular localization. Donut chart showing the major DeepLoc2.0 predictions of zebrafish proteome. The number of proteins (**B**) and protein-coding genes (**B’**) are shown. (**C**) Venn diagram comparing the number of genes with existing extracellular annotation records and DeepLoc 2.0-based ‘Extracellular’ predictions. (**D**) Venn diagram comparing genes with existing cell membrane localization annotation records and DeepLoc-based ‘cell membrane’ predictions.

Next, we manually assessed how reliable are DeepLoc 2.0 predictions for a subset of proteins. DeepLoc 2.0 successfully predicted the most abundant secreted proteins comprising of pancreatic enzymes (Prss, Cela, Cpa, Amy2a family) as extracellular proteins (coded by 24 genes)^30^ (**Supplementary Table 2**). DeepLoc 2.0 prediction efficiency was also high for the Wnt family of secreted proteins (coded by 24 genes). Next, we assessed the DeepLoc 2.0 predictions for the Fgf protein family, which includes signal peptide-less secreted (Fgf9, Fgf16, Fgf20) and non-secreted Fgf11 subfamily^31^. DeepLoc 2.0 successfully predicted 17 genes coding for canonical signal peptide-containing extracellular Fgf proteins (Fgf 3-8,10, 17-19, 21-23) and all 7 of the intracellular Fgf11 subfamily proteins as non-extracellular proteins. However, when classifying signal peptide-less extracellular Fgf’s, DeepLoc 2.0 only classified Fgf9 as extracellular and misclassified Fgf16, Fgf20a and Fgf20b as cytoplasmic proteins (**Supplementary Table 2**). Hence, although DeepLoc 2.0 is highly reliable in predicting zebrafish secretome, manual curation is required.

### Curation of zebrafish secretome and membrane receptome

To create a list of zebrafish secretome and receptome, we first combined our DeepLoc 2.0 predictions with annotations from other existing records (UniprotKB CC, GO terms, and Matrisome)^20,26,27^. In addition, we added the zebrafish orthologues of mammalian secretome- and receptor-coding genes based on Cell-Cell interaction (baderlab.org/CellCellInteractions) and CellTalkDB databases (**Figure 2A**)^32,33^ and performed manual curation (as detailed in Material sections). Our curated secretome (ligands) and cell membrane receptome database consist of 2788 (10.5%) and 2343 (8.8%) of protein-coding genes respectively (**Figure 2B &Supplementary Table 3**). In comparison to zebrafish matrisome dataset, our curated ligand lists contain 1848 additional genes (1566 DeepLoc 2 predictions and 282 manually curated extracellular-coding genes) (**Figure 2C**).

**Figure 2.**
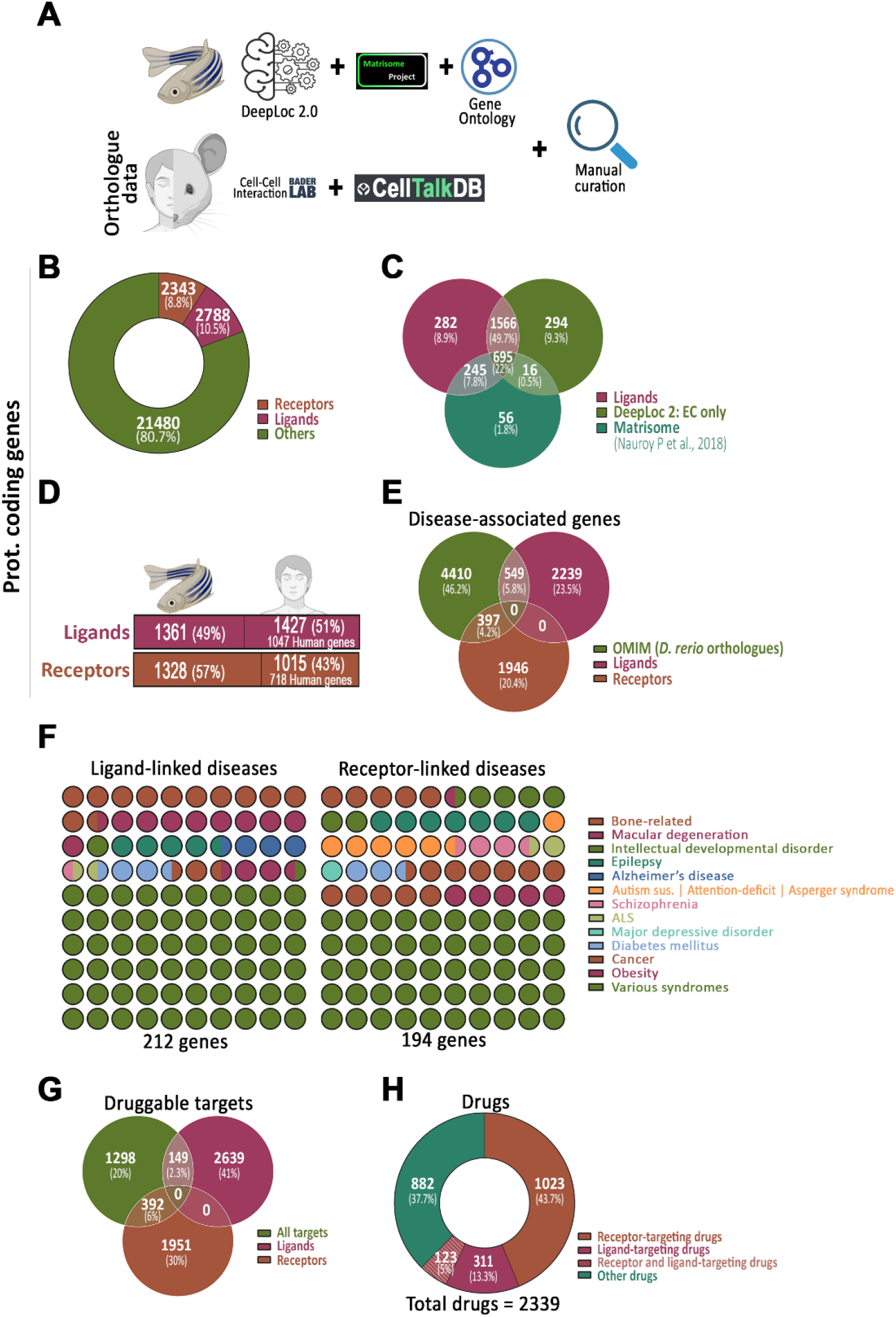
Curated zebrafish ligands and membrane receptors. (**A**) Scheme describing the steps taken for the creation of zebrafish ligands and membrane receptors list. (**B**) Donut chart showing the number and percentage of zebrafish genes coding for curated receptor- and ligand-coding genes. (**C**) Venn diagram comparing curated ligands with DeepLoc2.0 extracellular predicted proteins and previously reported Matrisome dataset. (**D**) Conservation between zebrafish and human ligand- and membrane receptor-coding genes based on ZFIN records. (**E**) Venn diagram comparing the number of curated ligand- and receptor-coding genes with zebrafish orthologues of human disease-linked genes in the OMIM database. (**F**) Dot plot showing the diseases associated with human orthologues of some of the zebrafish ligands and membrane receptors. (**G**) Venn diagram showing the number and percentage of zebrafish membrane receptors and ligands targetable by drugs. (**H**) Donut chart showing the number and percentage of drugs that can potentially target zebrafish receptors and/or ligands.

### Disease-associated secretome and membrane receptome

Next, we explored the identity of human diseases in which zebrafish models can be used for understanding disease etiology and/or in drug screening. We cross-referenced our ligand and membrane receptor list with the human orthologue information available in ZFIN database^34^. We identified 51% and 43% of the ligand- and cell membrane receptor-coding genes having human orthologue gene records (**Figure 2D & Supplementary Table 4**). Next, we cross-referenced our dataset with the zebrafish orthologues of human disease-associated genes in Online Mendelian Inheritance in Man (omim.org) database. We identified 549 ligand- and 397 receptor-coding genes containing records of human orthologue genes that are disease-associated (**Figure 2E**). Some of the major diseases associated with ligands and cell membrane receptors are related to bone diseases, macular degeneration, diabetes mellitus, and cancer **(Figure 2F)**. Nevertheless, several of the ligands and receptors were also associated with numerous syndromes (e.g., Ehlers-Danlos syndrome, Loeys-Dietz syndrome and Long QT syndrome) (**Figure 2F)**.

Next, we explored the drug-targetable status of the zebrafish secretome and receptome. We cross-referenced our ligand and receptome list with DrugCentral database that contains an up-to-date list of drugs and their targets^35^. We identified 149 ligands and 392 receptors that can be potentially targeted by drugs (**Figure 2G & Supplementary Table 5**). With respect to the choice of drugs, 311 (13%) and 1023 (44%) of the drugs can potentially target zebrafish ligands and cell membrane receptors respectively (**Figure 2H**).

### *DanioTalk* - putative *secretome-membrane receptome interactome in zebrafish*

Next, to map the ligand-receptome interactions, we identified all pairwise interactions between our ligand and receptome proteins in the STRING (v11.5) physical protein-protein interaction records of zebrafish (**Figure 3A**), a subset of which is shown in (**Figure 3A’**)^36^. The experimental score in STRING database predicts the confidence in the physical proximity of two proteins. Ligand-receptor pairs with scores with high-confidence scores are very likely to physically interact and result in associated downstream signaling activities. Our interactome database contains data for 370 ligands and 342 receptors at high confidence (**Figure 3B**). As we observed highly variable STRING physical interaction scores for experimentally validated orthologous human ligand-receptor pairs, we also incorporated medium-ranked pairs and orthologous human ligand-receptors pairs available in protein-protein interaction database IID (**Supplementary Figure S2**)^37^. This resulted in 10,446 ligand-receptor pairs (**Figure 3C, 3C’ & Supplementary Table 6**). Based on our analysis, at least 1681 ligands and 1455 receptors lack a high-ranking interacting receptor or ligand, respectively (**Figure 3B**).

**Figure 3.**
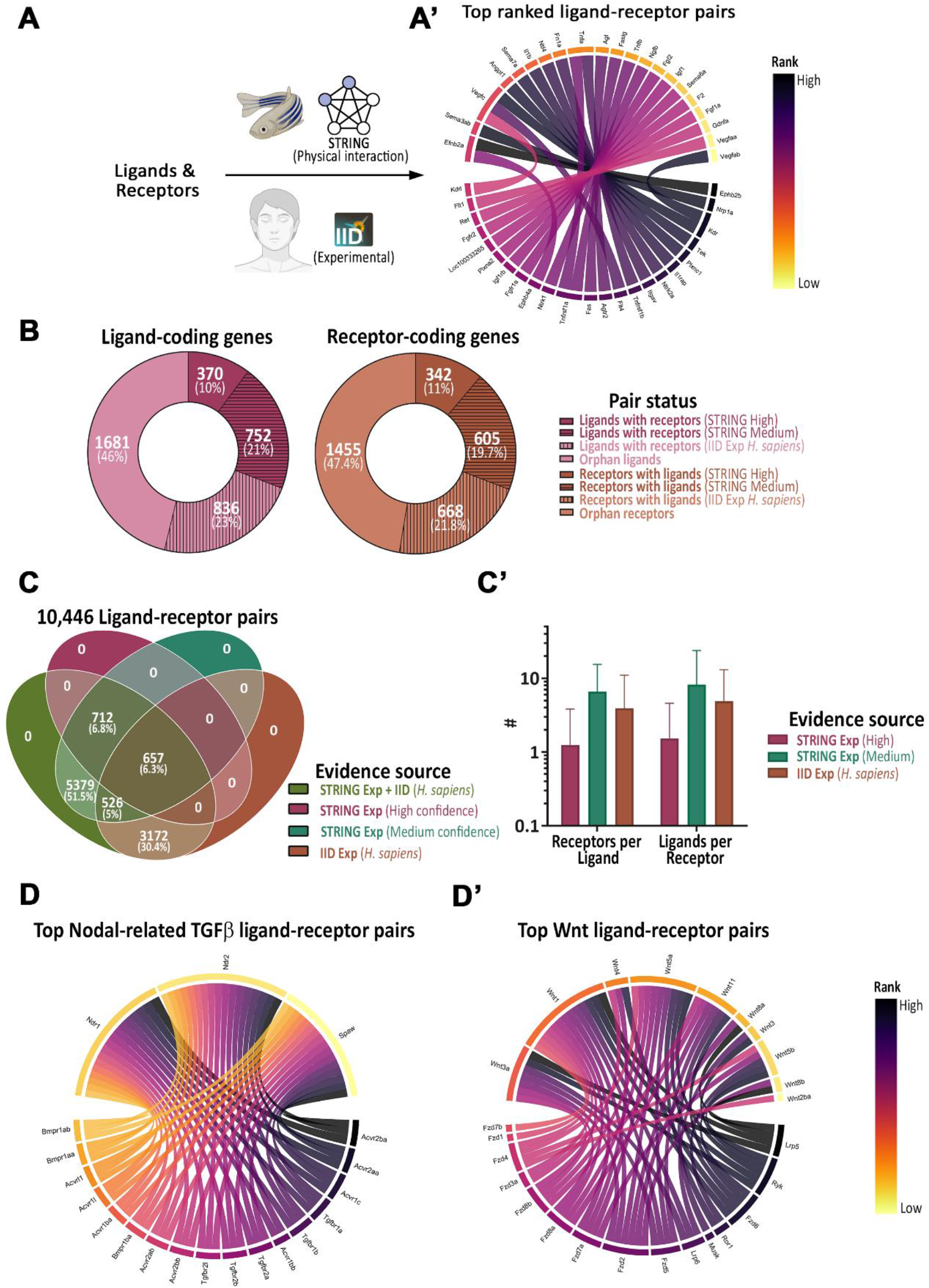
Zebrafish Ligand-Receptor Interactome. (**A**) Scheme describing steps undertaken to map the zebrafish ligands and membrane receptors interactome (**A**). Circle plot showing the top 25 ranked ligand-receptor pairs (**A’**). (**B**) Orphan status of ligands and membrane receptors. Donut chart showing the number and percentage of genes coding for ligands and receptors with or without potential receptors and ligands respectively. (**C**) Zebrafish ligand-receptor interactome stats. Venn diagram comparing ligand-receptor pairs across different ranking categories (**C**). Graph showing # of putative receptors/ligand and ligands/receptor (**C’**). (**D**) Receptors for Nodal-related TGF-beta ligands and Wnt ligands. Circle plot showing the top ranked receptors for Ndr2, Ndr1 and Spaw ligands (**D**) and Wnt ligands (**D’**).

We next probed the accuracy of our ligand-receptor interactome database. We focused on the Nodal-related TGFβ ligands and WNT ligands, key regulators of development^4,38,39^. Our interaction database identified all the previously reported receptors for Ndr2/Cyclops, Ndr1/Squint and Spaw/Southpaw ligands namely Acvr1b/Alk4 (rank 6 and 13), Acvr2a (rank 2 and 11), Acvr2b (rank 1 and 10) respectively (**Figure 3D &Supplementary Table 7**). We also noticed Acvr1c, Tgfbr1 and Tgfbr2 as top ranked receptors. Next, we probed Wnt signaling ligand-receptor interactome. Our interaction database identified all the Wnt receptors (Lrp5, Ryk, Fzd, Ror1 and Lrp6) that can interact with the major Wnt ligands (**Figure 3D’ & Supplementary Table 7**). Altogether, *DanioTalk* is able to accurately identify major ligand-receptor interactions in zebrafish.

### Zebrafish brain synaptic secretome-membrane receptome interactome

Next, we tested the application of *DanioTalk* on existing datasets. We first focused on the zebrafish adult brain synaptosomal and postsynaptic density (PSD) proteome dataset (**Figure 4A**)^40^. We identified protein products of 147 ligand- and 182 receptor-coding genes in the synaptosomal fraction. While in the PSD fraction, we identified protein products from 74 ligand- and 86 receptor-coding genes (**Supplementary Figure S3 & Supplementary Table 8**). The top-ranked ligand-receptor interacting pairs between synaptosome ligands and PSD receptors were Agrn-Ptprsa, Cntn2-Cntnap1/Cntn1a/Cntn1b, and Tncb/Adgrl1a (**Figure 4B**) (**Supplementary Table 9**). The top-ranked PSD ligand-synaptosome receptor pairs were Tncb-Itgav/Cntn1b and Cntn2-Nrcama/Cntnap1/Cntn1a (**Figure 4B’**) (**Supplementary Table 9**).

**Figure 4.**
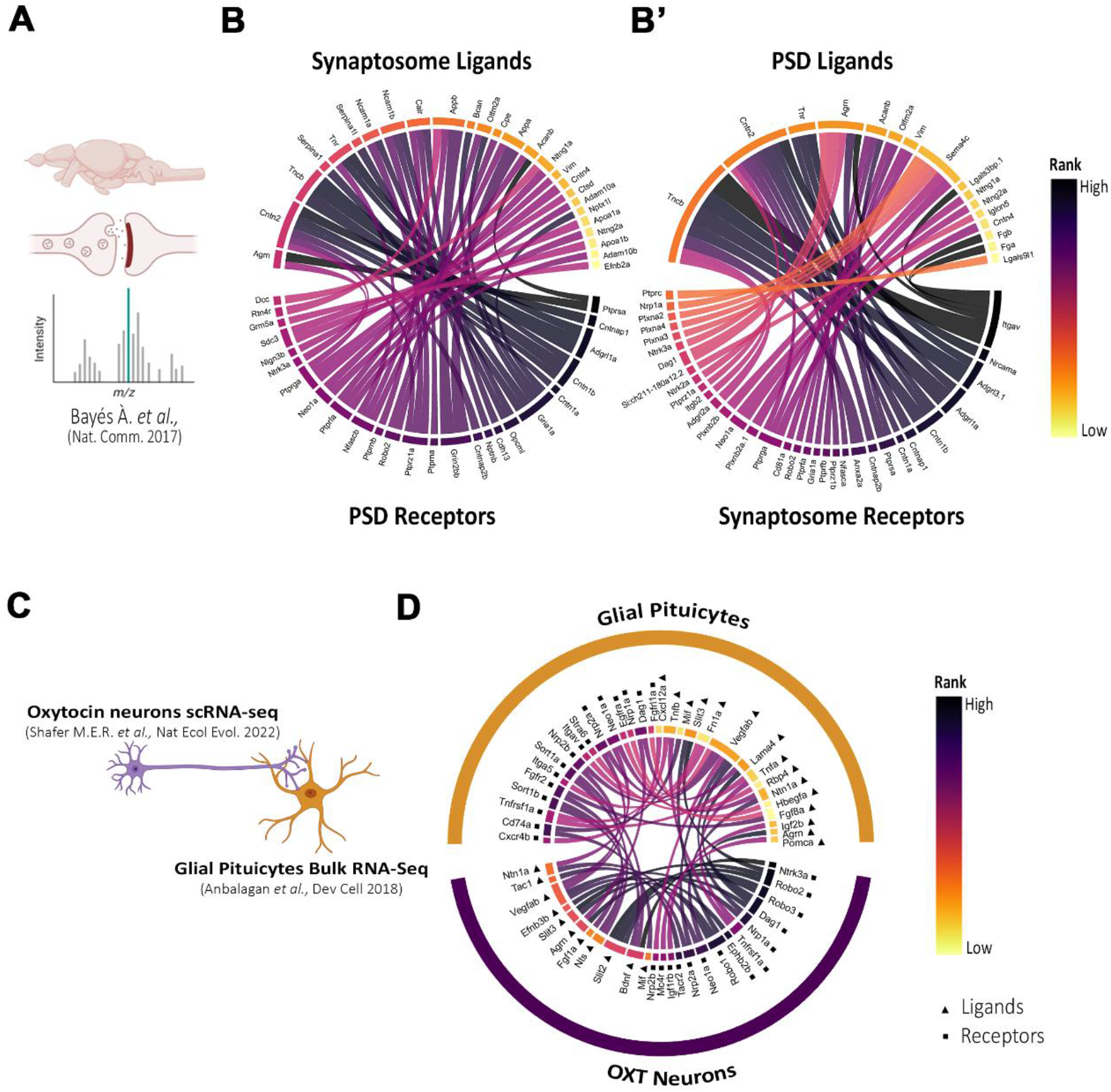
Application of DanioTalk. **(A)** Scheme showing synaptosome-PSD from adult zebrafish brain. **(B)** Circle plot showing the top-ranked ligand-receptor interactions between the synaptosome and PSD (**B** and **B’**). Interactions were ranked based on quasi-percentile of peptide expression and score >400 (Medium confidence). **(C)** Scheme showing axo-glial interaction between OXT neuron and glial pituicytes. **(D)** Circle plot showing top-ranked ligand-receptor interactions between OXT neurons and glial pituicytes. Interactions were ranked based on quasi-percentile of gene expression and score >900 (Highest confidence).

### Zebrafish Oxytocin Neuron-Glial Pituicytes Axo-Glial Interactome

Axo-glial interactions contribute to synaptic morphogenesis and plasticity^41,42^. However, the identify of all the axo-glial ligand-receptor interactions of neurohypophysis (NH), a major neuroendocrine interface has not been addressed^43,44^. The NH-projecting axonal tracts, NH axo-glial cellular components and their ultrastructure are conserved in zebrafish^45–48^. We aimed to identify the axo-glial interactome of Oxytocin (OXT) neurons and glial pituicytes. Due to the lack of OXT neuronal axonal transcriptome, we used the OXT neuron scRNA-seq data and glial pituicyte bulk RNA-seq data (**Figure 4C**)^45,49^. Using the differentially expressed list of genes from these datasets, we identified only 3 ligand-receptor interactions between OXT neurons and glial pituicytes at high confidence (Nts-Sort1a, Bdnf-Ngrfb/Sort1a) (**Supplementary Figure S3C-E & Supplementary Table 10)**. To incorporate all the genes expressed in OXT neurons, we used the pseudo-bulk expression data of OXT neuron enriched cluster. This led to the identification of 259 ligands and 209 receptors in OXT neurons. Whereas in glial pituicytes transcriptome, we identified 325 ligands and 174 receptors with average read counts >50 (**Supplementary Figure S3F-G & Supplementary Table 11**). Analyzing the ligand-receptor pairs in this dataset led to identification of 37 OXT-neuron-glial pituicyte ligand-receptor interactions. The top ranked pituicyte ligand-OXT neuron receptor interactions were Slit3-Robo2/Robo1, Lama4-Dag1 and Vegfab-Nrp1a. While the top ranked OXT neuron ligand-pituicyte receptors were Mif-Cd74a, Nts-Sort1a/Sort1b, and Bdnf-Sort1a/Sort1b (**Figure 4D & Supplementary Table 11**). We also observed several autocrine signaling interactions in glial pituicytes (Cxcl12a-Cxcr4b, Tnfb-Tnfrsf1a, and Fgf8a/Fgf10a-Fgfr2) and in OXT neurons (Slit3/Slit2-Robo2/Robo1/Robo3, Bdnf-Ntrk3a/Sort1a, and Agrn-Dag1) (**Figure 4D & Supplemental Table 11**).

## Discussion

Zebrafish single-cell RNAseq datasets are used to explore the developmental ligand-receptor interactome^23–25,50^. However, a bottleneck in using such tools is in the dependence on incomplete orthologous mammalian ligand-receptor classifications and mammalian ligand-receptor interactome records^51,52^. Thus, tools to study zebrafish secretome, receptome and interactome independent of orthologue records are essential to help explore ligand-receptor interactions in zebrafish datasets.

In the lack of experimental records, recent studies reported the relatively high success of machine learning-based tools in predicting protein structures and cellular localizations^28,53,54^. Here, we applied a deep learning algorithm to predict the cellular localization of the zebrafish proteome and created a curated repository of the zebrafish secretome and membrane receptome. This allows the inclusion of orphan zebrafish secretome or membranal receptor proteins that lack cellular localization or ortholog records. Also, our database of secretome and receptome-linked diseases will aid researchers interested in developing zebrafish models of human disease.

Generally, creating disease models of zebrafish may require the creation of mutations in paralogous genes^55,56^. Our results suggest that this is not the case for all diseases. For instance, secreted factor-coding genes such as *col13a1, cpe, dll4* and *tgfb2* can be potentially used to model congenital myasthenic syndrome 19, BDV syndrome, Adam-Oliver syndrome and Loeys-Dietz syndrome 4 respectively^57–60^. Whereas receptor coding genes such as *gpc4* and *ccln4* can be potentially used for studying Keipert syndrome and Raynaud syndrome respectively^61,62^.

One of the strengths of research on larval zebrafish is in the pharmacological perturbation experiments which we have also taken advantage in the past^45^. Some subscription-based tools built for mammalian model organisms can list drugs targeting mammalian proteins^63,64^. Our *DanioTalk* repository and output files contain similar information on drugs that can likely target the zebrafish secretome and receptome.

The potential of *DanioTalk* was revealed in our analysis of the existing dataset on adult zebrafish brain synaptosome-PSD proteomes^40^. Several top ligand-receptor pairs were previously studied for their synaptic role in other model organisms. For example, in mice CNTNAP1 (CASPR) is well known for its role synaptic plasticity ^65^ and Contactin has been shown to promote synaptic Cntnap1 surface localization via *Cis*-interaction^65^. But the fact that Cntn2 and Cntnap1 are enriched in synaptosome and PSD fractions suggests an experimental artefact or additional trans-synaptic roles for the Cntn2-Cntnap1 interaction^66,67^. In addition, another top-ranking ligand-receptor pair was Tenascin (Tncb)-Adhesion G Protein-Coupled Receptor L1 (Adgrl1a). Both TNC and ADGRL1 regulate synaptic development and function, but the role of TNC-ADGRL1 interaction has not be addressed yet^68–71^.

The presence of several conserved ligand-receptor pairs in the zebrafish synapses further stresses the importance of zebrafish in understanding the role of ligand-receptor in synapse biology and in drugs targeting synapses^14,72,73^. Although our synaptosome and PSD ligand-receptor pairs lack spatiotemporal accuracy, *DanioTalk* can be further combined with zebrafish single-cell brain transcriptome records to identify synaptosome-PSD ligand-receptor interactions for neuronal cell types of interest^51,74^.

In the absence of cell-specific proteome data, RNA-seq data in combination with protein interaction records are largely used to explore ligand-receptor-based cellular interactions^75,76^. However, such data were not explored to analyze axo-glial ligand-receptor interactions in neurohypophysis (NH), a major neuroendocrine interface critical in maintaining water balance and reproductive functions^45,77,78^. Except for few molecular players, the axo-glial interactions in NH were largely unknown^77,79–81^.

Our analysis identified the Slit3-Robo2 glial pituicyte-OXT neuron interaction, which has previously been reported to promote synaptic OXT levels^82^. However, we also observed autocrine Slit-Robo interactions in OXT neurons. Slit-Robo autocrine signaling in mice motor neurons has been shown to promote axon fasciculation; however, such defects were not reported for NH-projecting OXT axons in zebrafish^82,83^.

Apart from glia to neuronal signaling, neuron to glial signaling are also major players in neuro-glial signalling^41^. We found that OXT-neuron derived Neurotensin (Nts) potentially signaling to glial pituicytes via Sortilin (Sort1b) receptor. In vertebrate brain, neurotensin has been shown to be localized in hypothalamic neurons and in pituitary^84–86^. Neurotensin can regulate intracellular calcium levels in *in vitro* astrocyte cultures from rat ventral tegmental area^87^. But the role of OXT-derived Neurotensin in glial pituicytes function or intracellular calcium is unknown^88^.

We also observed potential Cxcl12a-Cxcr4b, and Fgf10a-Fgfr2-based autocrine interactions in glial pituicytes^89^. CXCL12-CXCR4 functions as a chemokine and neurotransmitter in the developing brain and as mitogen in brain tumors^90–93^. Although CXCL12-CXCR4 interaction is known to modulate NH-projecting vasopressin neuron firing pattern, the role of CXCL12-CXCR4 interaction in glial pituicytes have not been reported yet^94,95^. We also observed Fgf10a-Fgfr2-based autocrine interactions in glial pituicytes, which suggests that Fgf10a could have additional autocrine functions in glial pituicytes such as reactivity and/or proliferation^96–100^

Overall, *DanioTalk* offers a comprehensive repository of zebrafish secretome, receptome, secretome-receptome interaction atlas, and a novel data exploratory tool for zebrafish-based researchers.

## Methods

### Zebrafish dataset

Zebrafish reference proteome (UP000000437) sequences along with GO-term and cellular localization annotations were downloaded from UniprotKB^27^. Gene symbols, orthologue gene symbols, aliases/synonyms, gene descriptions, and ZFIN IDs were downloaded from the ZFIN database^34^. The extracellular protein-coding gene was identified based on the annotation terms ‘extracellular’ or ‘secreted’ or ‘ligands’. Cell membrane protein-coding genes were identified based on the terms “plasma membrane”.

### Subcellular localization prediction

Zebrafish reference proteome cellular localization was predicted using DeepLoc 2.0 with default parameters^28^.

### Curated database of the zebrafish secretome and membrane receptome

Human receptors and ligands were obtained from the Cell-Cell Interaction Database (Gary Bader group, Univ. of Toronto) and the CellTalkDB receptor database^32,33^. In addition human ligands were obtained from MatrisomeDB^101^. The initial list of zebrafish ligands was obtained by combining i) zebrafish matrisome, ii) zebrafish orthologues from the human ligand database (see above) and iii) extracellular proteins in DeepLoc 2.0-based extracellular protein coding genes. Initial zebrafish membrane-localized receptors list were obtained by combining zebrafish orthologues of the human receptor database (see above), and DeepLoc 2.0-based membranal protein-coding genes. The list was manually curated based on gene descriptions and localization records of human orthologues available in the Compartments database and the Human Protein Atlas^30,102^.

### Ligand-Receptor interactome database

The ligand-receptor interaction database was generated based on STRING physical interaction database for zebrafish (v11.5, Ref) and zebrafish orthologues of human protein interaction data available in the IID database (version 2021-05) ^36,37^. Pairs were ranked according to the STRING physical interaction score (Highest ≥ 900, High ≥ 700, Medium ≥ 400). Orthologous proteins with experimentally validated interactions in the IID database (*H. sapiens*) database were given a score of 500 for ranking purposes.

### Application of *DanioTalk* on zebrafish datasets

Zebrafish synapse and PSD proteome datasets were extracted from Supplementary Tables in^40^. For peptide expression, the average peptide count for synaptosome and post synaptic density was utilized. Zebrafish OXT neuron marker genes and pseudo-bulk expression were downloaded from Supplemental_data/3-marker_gene_lists/Drerio_zebrafish.markers.sub.csv (Neuronal_04_2 cluster, avg. Log2FC > 0.25, p-adj. < 0.05) and Supplemental_data/4-pseudobulk_expression/Subclusters-Danio_rerio.csv (Neuronal_04_2 cluster, expression >0.05) respectively^49^. Glial pituicytes genes were downloaded from the supplemental table 1 (For differentially expressed genes: Log2FC > 1, p-adj. < 0.05, average AMCA+ read count ≥ 50 and for all expressed genes average: AMCA+ read count ≥ 50)^45^. Gene symbols were updated based on ZFIN alias database. The ligand and receptors were compared and ranked based on *DanioTalk* interaction database and plotted using Circlize R package^103,104^.

## Data availability

The scripts used in this manuscript are available on GitHub at https://github.com/DanioTalk.

## Authors’ contributions

S.A. obtained funding, conceived, and designed the study. A.Z. contributed to study design and performed DeepLoc 2.0 predictions. M.C. wrote all the scripts and performed all the analysis. S.A. performed manual curation of ligands and receptors, wrote the manuscript and prepared figures. All authors read and approved of the manuscript.

## Acknowledgments

We thank Emilia Wysocka, Iwona Kanonik-Jędrzejak and Arleta Kucz for technical and administrative support. We thank Dr. Damian Szklarczyk (Swiss Institute of Bioinformatics, Zurich) for discussion and advice. We thank the BioRender.com team.

## References

1. Blockus, H., and Chédotal, A. (2016). Slit-Robo signaling. Development 143, 3037–3044. 10.1242/dev.132829.

2. Dufour, S., Quérat, B., Tostivint, H., Pasqualini, C., Vaudry, H., and Rousseau, K. (2020). Origin and Evolution of the Neuroendocrine Control of Reproduction in Vertebrates, With Special Focus on Genome and Gene Duplications. Physiological Reviews 100, 869–943. 10.1152/physrev.00009.2019.

3. Pires-daSilva, A., and Sommer, R.J. (2003). The evolution of signalling pathways in animal development. Nat Rev Genet 4, 39–49. 10.1038/nrg977.

4. Steinhart, Z., and Angers, S. (2018). Wnt signaling in development and tissue homeostasis. Development 145, dev146589. 10.1242/dev.146589.

5. Kowalczyk, W., Romanelli, L., Atkins, M., Hillen, H., Bravo González-Blas, C., Jacobs, J., Xie, J., Soheily, S., Verboven, E., Moya, I.M., et al. (2022). Hippo signaling instructs ectopic but not normal organ growth. Science 378, eabg3679. 10.1126/science.abg3679.

6. Nisar, S., Bhat, A.A., Masoodi, T., Hashem, S., Akhtar, S., Ali, T.A., Amjad, S., Chawla, S., Bagga, P., Frenneaux, M.P., et al. (2022). Genetics of glutamate and its receptors in autism spectrum disorder. Mol Psychiatry 27, 2380–2392. 10.1038/s41380-022-01506-w.

7. Straussman, R., Morikawa, T., Shee, K., Barzily-Rokni, M., Qian, Z.R., Du, J., Davis, A., Mongare, M.M., Gould, J., Frederick, D.T., et al. (2012). Tumour micro-environment elicits innate resistance to RAF inhibitors through HGF secretion. Nature 487, 500–504. 10.1038/nature11183.

8. Attwood, M.M., Jonsson, J., Rask-Andersen, M., and Schiöth, H.B. (2020). Soluble ligands as drug targets. Nat Rev Drug Discov 19, 695–710. 10.1038/s41573-020-0078-4.

9. Hauser, A.S., Attwood, M.M., Rask-Andersen, M., Schiöth, H.B., and Gloriam, D.E. (2017). Trends in GPCR drug discovery: new agents, targets and indications. Nat Rev Drug Discov 16, 829–842. 10.1038/nrd.2017.178.

10. Congreve, M., de Graaf, C., Swain, N.A., and Tate, C.G. (2020). Impact of GPCR Structures on Drug Discovery. Cell 181, 81–91. 10.1016/j.cell.2020.03.003.

11. Lees, J.A., Dias, J.M., and Han, S. (2021). Applications of Cryo-EM in small molecule and biologics drug design. Biochem Soc Trans 49, 2627–2638. 10.1042/BST20210444.

12. Tunyasuvunakool, K., Adler, J., Wu, Z., Green, T., Zielinski, M., Žídek, A., Bridgland, A., Cowie, A., Meyer, C., Laydon, A., et al. (2021). Highly accurate protein structure prediction for the human proteome. Nature 596, 590–596. 10.1038/s41586-021-03828-1.

13. Cagan, R.L., Zon, L.I., and White, R.M. (2019). Modeling Cancer with Flies and Fish. Dev Cell 49, 317–324. 10.1016/j.devcel.2019.04.013.

14. Hoffman, E.J., Turner, K.J., Fernandez, J.M., Cifuentes, D., Ghosh, M., Ijaz, S., Jain, R.A., Kubo, F., Bill, B.R., Baier, H., et al. (2016). Estrogens Suppress a Behavioral Phenotype in Zebrafish Mutants of the Autism Risk Gene, CNTNAP2. Neuron 89, 725–733. 10.1016/j.neuron.2015.12.039.

15. Patton, E.E., Zon, L.I., and Langenau, D.M. (2021). Zebrafish disease models in drug discovery: from preclinical modelling to clinical trials. Nat Rev Drug Discov 20, 611–628. 10.1038/s41573-021-00210-8.

16. Ségalat, L. (2007). Invertebrate animal models of diseases as screening tools in drug discovery. ACS Chem Biol 2, 231–236. 10.1021/cb700009m.

17. Csályi, K., Fazekas, D., Kadlecsik, T., Türei, D., Gul, L., Horváth, B., Módos, D., Demeter, A., Pápai, N., Lenti, K., et al. (2016). SignaFish: A Zebrafish-Specific Signaling Pathway Resource. Zebrafish 13, 541–544. 10.1089/zeb.2016.1277.

18. Goldberg, T., Hecht, M., Hamp, T., Karl, T., Yachdav, G., Ahmed, N., Altermann, U., Angerer, P., Ansorge, S., Balasz, K., et al. (2014). LocTree3 prediction of localization. Nucleic Acids Res 42, W350–355. 10.1093/nar/gku396.

19. Klee, E.W. (2008). The zebrafish secretome. Zebrafish 5, 131–138. 10.1089/zeb.2008.0529.

20. Nauroy, P., Hughes, S., Naba, A., and Ruggiero, F. (2018). The in-silico zebrafish matrisome: A new tool to study extracellular matrix gene and protein functions. Matrix Biol 65, 5–13. 10.1016/j.matbio.2017.07.001.

21. Dimitrov, D., Türei, D., Garrido-Rodriguez, M., Burmedi, P.L., Nagai, J.S., Boys, C., Ramirez Flores, R.O., Kim, H., Szalai, B., Costa, I.G., et al. (2022). Comparison of methods and resources for cell-cell communication inference from single-cell RNA-Seq data. Nat Commun 13, 3224. 10.1038/s41467-022-30755-0.

22. Schier, A.F. (2020). Single-cell biology: beyond the sum of its parts. Nat Methods 17, 17–20. 10.1038/s41592-019-0693-3.

23. Campbell, N.R., Rao, A., Hunter, M.V., Sznurkowska, M.K., Briker, L., Zhang, M., Baron, M., Heilmann, S., Deforet, M., Kenny, C., et al. (2021). Cooperation between melanoma cell states promotes metastasis through heterotypic cluster formation. Developmental Cell 56, 2808–2825.e10. 10.1016/j.devcel.2021.08.018.

24. Holler, K., Neuschulz, A., Drewe-Boß, P., Mintcheva, J., Spanjaard, B., Arsiè, R., Ohler, U., Landthaler, M., and Junker, J.P. (2021). Spatio-temporal mRNA tracking in the early zebrafish embryo. Nat Commun 12, 3358. 10.1038/s41467-021-23834-1.

25. Hu, B., Lelek, S., Spanjaard, B., El-Sammak, H., Simões, M.G., Mintcheva, J., Aliee, H., Schäfer, R., Meyer, A.M., Theis, F., et al. (2022). Origin and function of activated fibroblast states during zebrafish heart regeneration. Nat Genet 54, 1227–1237. 10.1038/s41588-022-01129-5.

26. Gene Ontology Consortium (2021). The Gene Ontology resource: enriching a GOld mine. Nucleic Acids Res 49, D325–D334. 10.1093/nar/gkaa1113.

27. The UniProt Consortium (2021). UniProt: the universal protein knowledgebase in 2021. Nucleic Acids Research 49, D480–D489. 10.1093/nar/gkaa1100.

28. Thumuluri, V., Almagro Armenteros, J.J., Johansen, A.R., Nielsen, H., and Winther, O. (2022). DeepLoc 2.0: multi-label subcellular localization prediction using protein language models. Nucleic Acids Res, gkac278. 10.1093/nar/gkac278.

29. Nevers, Y., Jones, T.E.M., Jyothi, D., Yates, B., Ferret, M., Portell-Silva, L., Codo, L., Cosentino, S., Marcet-Houben, M., Vlasova, A., et al. (2022). The Quest for Orthologs orthology benchmark service in 2022. Nucleic Acids Research 50, W623–W632. 10.1093/nar/gkac330.

30. Uhlén, M., Fagerberg, L., Hallström, B.M., Lindskog, C., Oksvold, P., Mardinoglu, A., Sivertsson, Å., Kampf, C., Sjöstedt, E., Asplund, A., et al. (2015). Proteomics. Tissue-based map of the human proteome. Science 347, 1260419. 10.1126/science.1260419.

31. Ornitz, D.M., and Itoh, N. (2015). The Fibroblast Growth Factor signaling pathway. Wiley Interdiscip Rev Dev Biol 4, 215–266. 10.1002/wdev.176.

32. Shao, X., Liao, J., Li, C., Lu, X., Cheng, J., and Fan, X. (2021). CellTalkDB: a manually curated database of ligand-receptor interactions in humans and mice. Brief Bioinform 22, bbaa269. 10.1093/bib/bbaa269.

33. Ximerakis, M., Lipnick, S.L., Innes, B.T., Simmons, S.K., Adiconis, X., Dionne, D., Mayweather, B.A., Nguyen, L., Niziolek, Z., Ozek, C., et al. (2019). Single-cell transcriptomic profiling of the aging mouse brain. Nat Neurosci 22, 1696–1708. 10.1038/s41593-019-0491-3.

34. Bradford, Y.M., Van Slyke, C.E., Ruzicka, L., Singer, A., Eagle, A., Fashena, D., Howe, D.G., Frazer, K., Martin, R., Paddock, H., et al. (2022). Zebrafish information network, the knowledgebase for Danio rerio research. Genetics 220, iyac016. 10.1093/genetics/iyac016.

35. Avram, S., Bologa, C.G., Holmes, J., Bocci, G., Wilson, T.B., Nguyen, D.-T., Curpan, R., Halip, L., Bora, A., Yang, J.J., et al. (2021). DrugCentral 2021 supports drug discovery and repositioning. Nucleic Acids Res 49, D1160–D1169. 10.1093/nar/gkaa997.

36. Szklarczyk, D., Gable, A.L., Nastou, K.C., Lyon, D., Kirsch, R., Pyysalo, S., Doncheva, N.T., Legeay, M., Fang, T., Bork, P., et al. (2021). The STRING database in 2021: customizable protein-protein networks, and functional characterization of user-uploaded gene/measurement sets. Nucleic Acids Res 49, D605–D612. 10.1093/nar/gkaa1074.

37. Kotlyar, M., Pastrello, C., Ahmed, Z., Chee, J., Varyova, Z., and Jurisica, I. (2022). IID 2021: towards context-specific protein interaction analyses by increased coverage, enhanced annotation and enrichment analysis. Nucleic Acids Res 50, D640–D647. 10.1093/nar/gkab1034.

38. Schier, A.F., and Shen, M.M. (2000). Nodal signalling in vertebrate development. Nature 403, 385–389. 10.1038/35000126.

39. Shen, M.M. (2007). Nodal signaling: developmental roles and regulation. Development 134, 1023–1034. 10.1242/dev.000166.

40. Bayés, À., Collins, M.O., Reig-Viader, R., Gou, G., Goulding, D., Izquierdo, A., Choudhary, J.S., Emes, R.D., and Grant, S.G.N. (2017). Evolution of complexity in the zebrafish synapse proteome. Nat Commun 8, 14613. 10.1038/ncomms14613.

41. Allen, N.J., and Eroglu, C. (2017). Cell Biology of Astrocyte-Synapse Interactions. Neuron 96, 697–708. 10.1016/j.neuron.2017.09.056.

42. Allen, N.J., and Lyons, D.A. (2018). Glia as architects of central nervous system formation and function. Science 362, 181–185. 10.1126/science.aat0473.

43. Pearson, C.A., and Placzek, M. (2013). Development of the medial hypothalamus: forming a functional hypothalamic-neurohypophyseal interface. Curr Top Dev Biol 106, 49–88. 10.1016/B978-0-12-416021-7.00002-X.

44. Wittkowski, W. (1998). Tanycytes and pituicytes: morphological and functional aspects of neuroglial interaction. Microsc Res Tech 41, 29–42. 10.1002/(SICI)1097-0029(19980401)41:1<29::AID-JEMT4>3.0.CO;2-P.

45. Anbalagan, S., Gordon, L., Blechman, J., Matsuoka, R.L., Rajamannar, P., Wircer, E., Biran, J., Reuveny, A., Leshkowitz, D., Stainier, D.Y.R., et al. (2018). Pituicyte Cues Regulate the Development of Permeable Neuro-Vascular Interfaces. Dev Cell 47, 711–726.e5. 10.1016/j.devcel.2018.10.017.

46. Grinevich, V., and Neumann, I.D. (2021). Brain oxytocin: how puzzle stones from animal studies translate into psychiatry. Mol Psychiatry 26, 265–279. 10.1038/s41380-020-0802-9.

47. Gutnick, A., Blechman, J., Kaslin, J., Herwig, L., Belting, H.-G., Affolter, M., Bonkowsky, J.L., and Levkowitz, G. (2011). The hypothalamic neuropeptide oxytocin is required for formation of the neurovascular interface of the pituitary. Dev Cell 21, 642–654. 10.1016/j.devcel.2011.09.004.

48. Herget, U., Gutierrez-Triana, J.A., Salazar Thula, O., Knerr, B., and Ryu, S. (2017). Single-Cell Reconstruction of Oxytocinergic Neurons Reveals Separate Hypophysiotropic and Encephalotropic Subtypes in Larval Zebrafish. eNeuro 4, ENEURO.0278-16.2016. 10.1523/ENEURO.0278-16.2016.

49. Shafer, M.E.R., Sawh, A.N., and Schier, A.F. (2022). Gene family evolution underlies cell-type diversification in the hypothalamus of teleosts. Nat Ecol Evol 6, 63–76. 10.1038/s41559-021-01580-3.

50. Herpelinck, T., Ory, L., Nasello, G., Barzegari, M., Bolander, J., Luyten, F.P., Tylzanowski, P., and Geris, L. (2022). An integrated single-cell atlas of the skeleton from development through adulthood. 2022.03.14.484345. 10.1101/2022.03.14.484345.

51. Farnsworth, D.R., Saunders, L.M., and Miller, A.C. (2020). A single-cell transcriptome atlas for zebrafish development. Dev Biol 459, 100–108. 10.1016/j.ydbio.2019.11.008.

52. Farrell, J.A., Wang, Y., Riesenfeld, S.J., Shekhar, K., Regev, A., and Schier, A.F. (2018). Single-cell reconstruction of developmental trajectories during zebrafish embryogenesis. Science 360, eaar3131. 10.1126/science.aar3131.

53. Jumper, J., Evans, R., Pritzel, A., Green, T., Figurnov, M., Ronneberger, O., Tunyasuvunakool, K., Bates, R., Žídek, A., Potapenko, A., et al. (2021). Highly accurate protein structure prediction with AlphaFold. Nature 596, 583–589. 10.1038/s41586-021-03819-2.

54. Lin, Z., Akin, H., Rao, R., Hie, B., Zhu, Z., Lu, W., Smetanin, N., Verkuil, R., Kabeli, O., Shmueli, Y., et al. (2022). Evolutionary-scale prediction of atomic level protein structure with a language model. 2022.07.20.500902. 10.1101/2022.07.20.500902.

55. Hinman, M.N., Richardson, J.I., Sockol, R.A., Aronson, E.D., Stednitz, S.J., Murray, K.N., Berglund, J.A., and Guillemin, K. (2021). Zebrafish mbnl mutants model physical and molecular phenotypes of myotonic dystrophy. Dis Model Mech 14, dmm045773. 10.1242/dmm.045773.

56. Howe, K., Clark, M.D., Torroja, C.F., Torrance, J., Berthelot, C., Muffato, M., Collins, J.E., Humphray, S., McLaren, K., Matthews, L., et al. (2013). The zebrafish reference genome sequence and its relationship to the human genome. Nature 496, 498–503. 10.1038/nature12111.

57. Bosch, E., Hebebrand, M., Popp, B., Penger, T., Behring, B., Cox, H., Towner, S., Kraus, C., Wilson, W.G., Khan, S., et al. (2021). BDV Syndrome: An Emerging Syndrome With Profound Obesity and Neurodevelopmental Delay Resembling Prader-Willi Syndrome. J Clin Endocrinol Metab 106, 3413–3427. 10.1210/clinem/dgab592.

58. Fontana, P., Genesio, R., Casertano, A., Cappuccio, G., Mormile, A., Nitsch, L., Iolascon, A., Andria, G., and Melis, D. (2014). Loeys-Dietz syndrome type 4, caused by chromothripsis, involving the TGFB2 gene. Gene 538, 69–73. 10.1016/j.gene.2014.01.017.

59. Logan, C.V., Cossins, J., Rodríguez Cruz, P.M., Parry, D.A., Maxwell, S., Martínez-Martínez, P., Riepsaame, J., Abdelhamed, Z.A., Lake, A.V.R., Moran, M., et al. (2015). Congenital Myasthenic Syndrome Type 19 Is Caused by Mutations in COL13A1, Encoding the Atypical Non-fibrillar Collagen Type XIII α1 Chain. Am J Hum Genet 97, 878–885. 10.1016/j.ajhg.2015.10.017.

60. Meester, J.A.N., Southgate, L., Stittrich, A.-B., Venselaar, H., Beekmans, S.J.A., den Hollander, N., Bijlsma, E.K., Helderman-van den Enden, A., Verheij, J.B.G.M., Glusman, G., et al. (2015). Heterozygous Loss-of-Function Mutations in DLL4 Cause Adams-Oliver Syndrome. Am J Hum Genet 97, 475–482. 10.1016/j.ajhg.2015.07.015.

61. Amor, D.J., Stephenson, S.E.M., Mustapha, M., Mensah, M.A., Ockeloen, C.W., Lee, W.S., Tankard, R.M., Phelan, D.G., Shinawi, M., de Brouwer, A.P.M., et al. (2019). Pathogenic Variants in GPC4 Cause Keipert Syndrome. Am J Hum Genet 104, 914–924. 10.1016/j.ajhg.2019.02.026.

62. Palmer, E.E., Stuhlmann, T., Weinert, S., Haan, E., Van Esch, H., Holvoet, M., Boyle, J., Leffler, M., Raynaud, M., Moraine, C., et al. (2018). De novo and inherited mutations in the X-linked gene CLCN4 are associated with syndromic intellectual disability and behavior and seizure disorders in males and females. Mol Psychiatry 23, 222–230. 10.1038/mp.2016.135.

63. Krämer, A., Green, J., Pollard, J., and Tugendreich, S. (2014). Causal analysis approaches in Ingenuity Pathway Analysis. Bioinformatics 30, 523–530. 10.1093/bioinformatics/btt703.

64. Stelzer, G., Rosen, N., Plaschkes, I., Zimmerman, S., Twik, M., Fishilevich, S., Stein, T.I., Nudel, R., Lieder, I., Mazor, Y., et al. (2016). The GeneCards Suite: From Gene Data Mining to Disease Genome Sequence Analyses. Current Protocols in Bioinformatics 54, 1.30.1–1.30.33. 10.1002/cpbi.5.

65. Murai, K.K., Misner, D., and Ranscht, B. (2002). Contactin Supports Synaptic Plasticity Associated with Hippocampal Long-Term Depression but Not Potentiation. Current Biology 12, 181–190. 10.1016/S0960-9822(02)00680-2.

66. Dubessy, A., Mazuir, E., Rappeneau, Q., Ou, S., Abi Ghanem, C., Piquand, K., Aigrot, M., Thétiot, M., Desmazières, A., Chan, E., et al. (2019). Role of a Contactin multi-molecular complex secreted by oligodendrocytes in nodal protein clustering in the CNS. Glia 67, 2248–2263. 10.1002/glia.23681.

67. Ruegg, M.A., Stoeckli, E.T., Kuhn, T.B., Heller, M., Zuellig, R., and Sonderegger, P. (1989). Purification of axonin-1, a protein that is secreted from axons during neurogenesis. EMBO J 8, 55–63. 10.1002/j.1460-2075.1989.tb03348.x.

68. Gurevicius, K., Kuang, F., Stoenica, L., Irintchev, A., Gureviciene, I., Dityatev, A., Schachner, M., and Tanila, H. (2009). Genetic ablation of tenascin-C expression leads to abnormal hippocampal CA1 structure and electrical activity in vivo. Hippocampus 19, 1232–1246. 10.1002/hipo.20585.

69. Vitobello, A., Mazel, B., Lelianova, V.G., Zangrandi, A., Petitto, E., Suckling, J., Salpietro, V., Meyer, R., Elbracht, M., Kurth, I., et al. (2022). ADGRL1 haploinsufficiency causes a variable spectrum of neurodevelopmental disorders in humans and alters synaptic activity and behavior in a mouse model. Am J Hum Genet 109, 1436–1457. 10.1016/j.ajhg.2022.06.011.

70. Li, J., Shalev-Benami, M., Sando, R., Jiang, X., Kibrom, A., Wang, J., Leon, K., Katanski, C., Nazarko, O., Lu, Y.C., et al. (2018). Structural Basis for Teneurin Function in Circuit-Wiring: A Toxin Motif at the Synapse. Cell 173, 735–748.e15. 10.1016/j.cell.2018.03.036.

71. Gottschling, C., Wegrzyn, D., Denecke, B., and Faissner, A. (2019). Elimination of the four extracellular matrix molecules tenascin-C, tenascin-R, brevican and neurocan alters the ratio of excitatory and inhibitory synapses. Sci Rep 9, 13939. 10.1038/s41598-019-50404-9.

72. Hutson, L.D., and Chien, C.-B. (2002). Wiring the zebrafish: axon guidance and synaptogenesis. Curr Opin Neurobiol 12, 87–92. 10.1016/s0959-4388(02)00294-5.

73. Oprişoreanu, A.-M., Smith, H.L., Krix, S., Chaytow, H., Carragher, N.O., Gillingwater, T.H., Becker, C.G., and Becker, T. (2021). Automated in vivo drug screen in zebrafish identifies synapse-stabilising drugs with relevance to spinal muscular atrophy. Dis Model Mech 14, dmm047761. 10.1242/dmm.047761.

74. Raj, B., Farrell, J.A., Liu, J., El Kholtei, J., Carte, A.N., Navajas Acedo, J., Du, L.Y., McKenna, A., Relić, θ., Leslie, J.M., et al. (2020). Emergence of Neuronal Diversity during Vertebrate Brain Development. Neuron 108, 1058–1074.e6. 10.1016/j.neuron.2020.09.023.

75. Efremova, M., Vento-Tormo, M., Teichmann, S.A., and Vento-Tormo, R. (2020). CellPhoneDB: inferring cell–cell communication from combined expression of multi-subunit ligand–receptor complexes. Nat Protoc 15, 1484–1506. 10.1038/s41596-020-0292-x.

76. Türei, D., Valdeolivas, A., Gul, L., Palacio-Escat, N., Klein, M., Ivanova, O., Ölbei, M., Gábor, A., Theis, F., Módos, D., et al. (2021). Integrated intra-and intercellular signaling knowledge for multicellular omics analysis. Molecular Systems Biology 17, e9923. 10.15252/msb.20209923.

77. Brown, C.H. (2016). Magnocellular Neurons and Posterior Pituitary Function. Compr Physiol 6, 1701–1741. 10.1002/cphy.c150053.

78. Romanov, R.A., Zeisel, A., Bakker, J., Girach, F., Hellysaz, A., Tomer, R., Alpár, A., Mulder, J., Clotman, F., Keimpema, E., et al. (2017). Molecular interrogation of hypothalamic organization reveals distinct dopamine neuronal subtypes. Nat Neurosci 20, 176–188. 10.1038/nn.4462.

79. Leng, G., Pineda, R., Sabatier, N., and Ludwig, M. (2015). 60 YEARS OF NEUROENDOCRINOLOGY: The posterior pituitary, from Geoffrey Harris to our present understanding. J Endocrinol 226, T173–185. 10.1530/JOE-15-0087.

80. Miyata, S. (2017). Advances in Understanding of Structural Reorganization in the Hypothalamic Neurosecretory System. Front Endocrinol (Lausanne) 8, 275. 10.3389/fendo.2017.00275.

81. Rosso, L., and Mienville, J.-M. (2009). Pituicyte modulation of neurohormone output. Glia 57, 235–243. 10.1002/glia.20760.

82. Anbalagan, S., Blechman, J., Gliksberg, M., Gordon, L., Rotkopf, R., Dadosh, T., Shimoni, E., and Levkowitz, G. (2019). Robo2 regulates synaptic oxytocin content by affecting actin dynamics. Elife 8, e45650. 10.7554/eLife.45650.

83. Jaworski, A., and Tessier-Lavigne, M. (2012). Autocrine/juxtaparacrine regulation of axon fasciculation by Slit-Robo signaling. Nat Neurosci 15, 367–369. 10.1038/nn.3037.

84. Goedert, M., Lightman, S.L., Mantyh, P.W., Hunt, S.P., and Emson, P.C. (1985). Neurotensin-like immunoreactivity and neurotensin receptors in the rat hypothalamus and in the neurointermediate lobe of the pituitary gland. Brain Res 358, 59–69. 10.1016/0006-8993(85)90948-5.

85. Batten, T.F., Marivoet, S., and Vandesande, F. (1987). Neurotensin-like immunoreactivity in the pituitary and hypothalamus of bony fishes. Peptides 8, 135–143. 10.1016/0196-9781(87)90177-x.

86. Muraki, K., Okahata, H., Nishi, Y., Usui, T., Yamada, H., Fujita, S., Miyachi, Y., Yanaihara, N., and Yajima, H. (1985). Distribution of neurotensin-like immunoreactivity in the hypothalamus, pituitary gland, and gastro-intestinal tract of rats. Acta Endocrinol (Copenh) 110, 1–5. 10.1530/acta.0.1100001.

87. Trudeau, L.E. (2000). Neurotensin regulates intracellular calcium in ventral tegmental area astrocytes: evidence for the involvement of multiple receptors. Neuroscience 97, 293–302. 10.1016/s0306-4522(99)00597-7.

88. Hatton, G.I., Bicknell, R.J., Hoyland, J., Bunting, R., and Mason, W.T. (1992). Arginine vasopressin mobilises intracellular calcium via V1-receptor activation in astrocytes (pituicytes) cultured from adult rat neural lobes. Brain Res 588, 75–83. 10.1016/0006-8993(92)91346-g.

89. Hickey, K.N., Grassi, S.M., Caplan, M.R., and Stabenfeldt, S.E. (2021). Stromal Cell-Derived Factor-1a Autocrine/Paracrine Signaling Contributes to Spatiotemporal Gradients in the Brain. Cell Mol Bioeng 14, 75–87. 10.1007/s12195-020-00643-y.

90. Barbero, S., Bajetto, A., Bonavia, R., Porcile, C., Piccioli, P., Pirani, P., Ravetti, J.L., Zona, G., Spaziante, R., Florio, T., et al. (2002). Expression of the chemokine receptor CXCR4 and its ligand stromal cell-derived factor 1 in human brain tumors and their involvement in glial proliferation in vitro. Ann N Y Acad Sci 973, 60–69. 10.1111/j.1749-6632.2002.tb04607.x.

91. Lazarini, F., Tham, T.N., Casanova, P., Arenzana-Seisdedos, F., and Dubois-Dalcq, M. (2003). Role of the alpha-chemokine stromal cell-derived factor (SDF-1) in the developing and mature central nervous system. Glia 42, 139–148. 10.1002/glia.10139.

92. Bhattacharyya, B.J., Banisadr, G., Jung, H., Ren, D., Cronshaw, D.G., Zou, Y., and Miller, R.J. (2008). The chemokine stromal cell-derived factor-1 regulates GABAergic inputs to neural progenitors in the postnatal dentate gyrus. J Neurosci 28, 6720–6730. 10.1523/JNEUROSCI.1677-08.2008.

93. Miyasaka, N., Knaut, H., and Yoshihara, Y. (2007). Cxcl12/Cxcr4 chemokine signaling is required for placode assembly and sensory axon pathfinding in the zebrafish olfactory system. Development 134, 2459–2468. 10.1242/dev.001958.

94. Callewaere, C., Banisadr, G., Desarménien, M.G., Mechighel, P., Kitabgi, P., Rostène, W.H., and Mélik Parsadaniantz, S. (2006). The chemokine SDF-1/CXCL12 modulates the firing pattern of vasopressin neurons and counteracts induced vasopressin release through CXCR4. Proc Natl Acad Sci U S A 103, 8221–8226. 10.1073/pnas.0602620103.

95. Callewaere, C., Fernette, B., Raison, D., Mechighel, P., Burlet, A., Calas, A., Kitabgi, P., Parsadaniantz, S.M., and Rostène, W. (2008). Cellular and subcellular evidence for neuronal interaction between the chemokine stromal cell-derived factor-1/CXCL 12 and vasopressin: regulation in the hypothalamo-neurohypophysial system of the Brattleboro rats. Endocrinology 149, 310–319. 10.1210/en.2007-1097.

96. Gómez-Pinilla, F., Vu, L., and Cotman, C.W. (1995). Regulation of astrocyte proliferation by FGF-2 and heparan sulfate in vivo. J Neurosci 15, 2021–2029. 10.1523/JNEUROSCI.15-03-02021.1995.

97. Goodman, T., Nayar, S.G., Clare, S., Mikolajczak, M., Rice, R., Mansour, S., Bellusci, S., and Hajihosseini, M.K. (2020). Fibroblast growth factor 10 is a negative regulator of postnatal neurogenesis in the mouse hypothalamus. Development 147, dev180950. 10.1242/dev.180950.

98. Haan, N., Goodman, T., Najdi-Samiei, A., Stratford, C.M., Rice, R., El Agha, E., Bellusci, S., and Hajihosseini, M.K. (2013). Fgf10-expressing tanycytes add new neurons to the appetite/energy-balance regulating centers of the postnatal and adult hypothalamus. J Neurosci 33, 6170–6180. 10.1523/JNEUROSCI.2437-12.2013.

99. Liu, F., Pogoda, H.-M., Pearson, C.A., Ohyama, K., Löhr, H., Hammerschmidt, M., and Placzek, M. (2013). Direct and indirect roles of Fgf3 and Fgf10 in innervation and vascularisation of the vertebrate hypothalamic neurohypophysis. Development 140, 1111–1122. 10.1242/dev.080226.

100. Zhang, X., Ibrahimi, O.A., Olsen, S.K., Umemori, H., Mohammadi, M., and Ornitz, D.M. (2006). Receptor Specificity of the Fibroblast Growth Factor Family: THE COMPLETE MAMMALIAN FGF FAMILY *. Journal of Biological Chemistry 281, 15694–15700. 10.1074/jbc.M601252200.

101. Shao, X., Taha, I.N., Clauser, K.R., Gao, Y. (Tom), and Naba, A. (2020). MatrisomeDB: the ECM-protein knowledge database. Nucleic Acids Research 48, D1136–D1144. 10.1093/nar/gkz849.

102. Binder, J.X., Pletscher-Frankild, S., Tsafou, K., Stolte, C., O’Donoghue, S.I., Schneider, R., and Jensen, L.J. (2014). COMPARTMENTS: unification and visualization of protein subcellular localization evidence. Database 2014, bau012. 10.1093/database/bau012.

103. Gu, Z., Gu, L., Eils, R., Schlesner, M., and Brors, B. (2014). circlize Implements and enhances circular visualization in R. Bioinformatics 30, 2811–2812. 10.1093/bioinformatics/btu393.

104. Team, R.C. (2013). R: A language and environment for statistical computing. R Foundation for Statistical Computing, Vienna, Austria. http://www.R-project.org/.

