## Supplemental figures for "A ligand-receptor interactome atlas of the zebrafish"

Supplementary Information

A

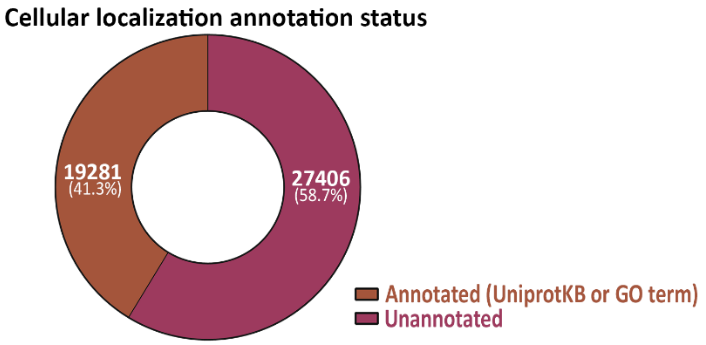


B

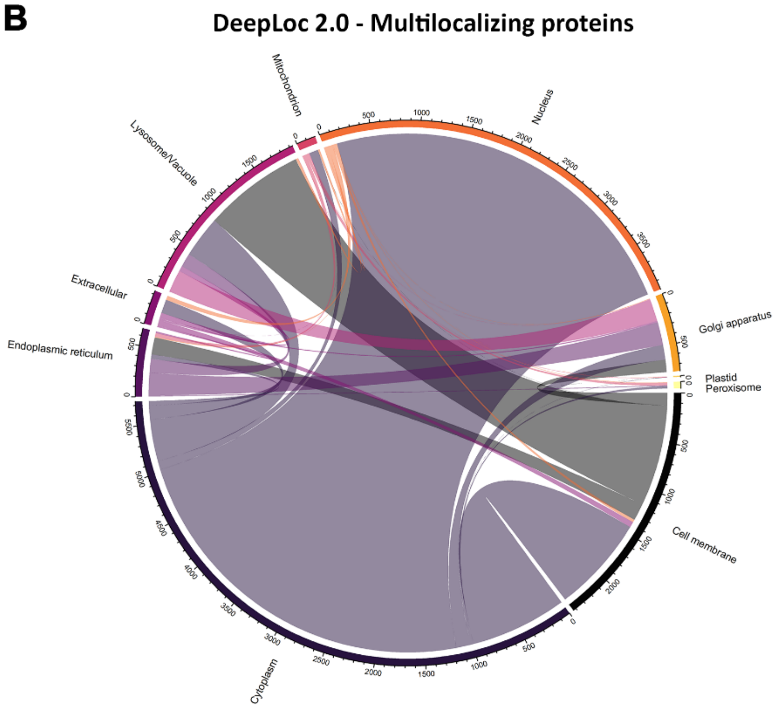


Supplementary Figure 1. Annotation status and DeepLoc 2.0 multilocalizing protein localizations

(A) Current status of the cellular localization annotation records of the reference proteome from zebrafish. Donut chart showing number of proteins with and without GO-term based or UniprotKB cellular localization records.

(B) Circle plot showing the number of proteins predicted to localize to multiple cellular regions and organelles.

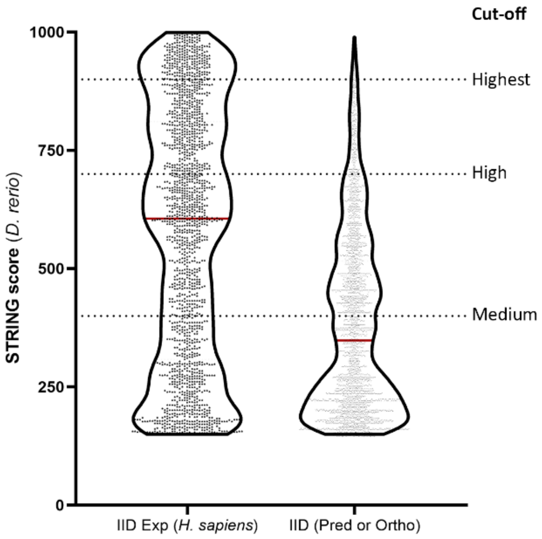


Supplementary Figure 2. Conservation of zebrafish ligand-receptor interactions.

Violin plot showing STRING database zebrafish ligand-receptor interactions scores to their human orthologue ligand-receptor records in IID database. 'Pred' denotes predicted interactions.

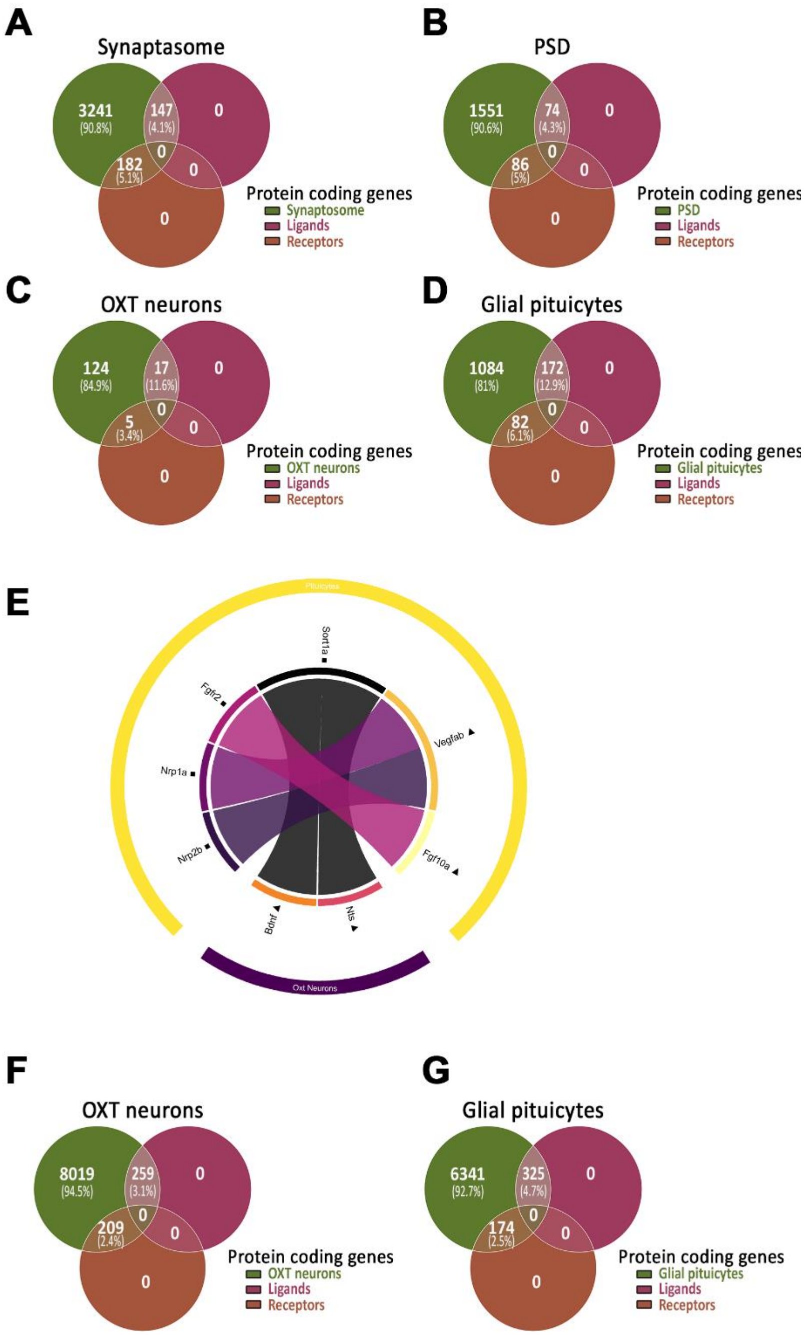


Supplementary Figure 3. Application of *DanioTalk*

(A) Venn diagram showing the number and percentage of genes coding for synaptosome-enriched ligands and receptors.

(B) Venn diagram showing the number and percentage of genes coding for PSD-enriched ligands and receptors.

(C) Venn diagram showing the number and percentage of ligand- and receptor-coding genes identified in OXT scRNA-seq differentially expressed dataset (log2FC>2, p-adj<0.05).

(D) Venn diagram showing the number and percentage of ligand- and receptor-coding genes identified in Glial pituicytes differentially expressed dataset (log2FC >2, p-adj<0.05, average AMCA+ read count >50).

(E) Circle plot showing top-ranked differentially expressed, ligand-receptor interactions between OXT neurons and glial pituicytes. Interactions were ranked based on gene expression and STRING physical interaction score (>900).

(F) Venn diagram showing the number and percentage of ligand- and receptor-coding genes identified in OXT pseudo-bulk RNA seq dataset (Expression >0.05).

(G) Venn diagram showing the number and percentage of ligand- and receptor-coding genes identified in Glial pituicytes differentially expressed dataset (Average AMCA+ read count >50).

Supplementary Tables

Supplementary table 1: Cellular localization predictions for zebrafish reference proteome.

Supplementary table 2: Cellular localization predictions for selected subset of zebrafish proteins

Supplementary table 3: Curated zebrafish secretome and membrane receptome

Supplementary table 4: List of zebrafish secretome and receptome coding genes with orthologues in OMIM database

Supplementary table 5: List of zebrafish secretome and receptome with orthologous proteins that are known drug targets as DrugCentralDB.

Supplementary table 6: List of zebrafish ligand-receptor pairs.

Supplementary table 7: List of ligand-receptors pairs in Nodal and Wnt signaling.

Supplementary table 8: List of ligand and receptor coding genes in Synaptosome and post-synaptic density.

Supplementary table 9: List of ligand-receptor pairs between Synaptosome and post-synaptic density.

Supplementary table 10: List of differentially expressed ligand and receptor coding genes and pairs in glial pituicytes and oxytocin neurons.

Supplementary table 11: List of all ligand and receptor coding genes and pairs between glial pituicytes and oxytocin neurons.
